# ConvNets Develop Organizational Principles of the Visual Cortex when using Ganglion Cell-Based Sampling

**DOI:** 10.1101/2021.11.02.466130

**Authors:** Danny da Costa, Rainer Goebel, Mario Senden

## Abstract

The distribution of retinal ganglion cells in primate visual systems portrays a densely distributed central region, with an incrementally decreasing cell density as the angle of visual eccentricity increases. This results in a non-uniform sampling of the retinal image that resembles a wheelbarrow distortion. We propose that this sampling gives rise to several organizational properties of the primate visual system, including cortical magnification, linear relationship between eccentricity and receptive field sizes, eccentricity-dependent drop-off in spatial-frequency preference, and radial bias. We test this hypothesis by training a convolutional neural network to classify the orientation of sine gratings and Gabor stimuli, resampled according to retinal ganglion cell distributions. Our simulations show that introducing this sampling step gives rise to the aforementioned organizational principles in convolutional layers while only minimally affecting their classification performance. This lends credence to the notion that the retinal ganglion cell distribution is an important factor for the emergence of these organizational principles in visual systems.

## 1 Introduction

A recent trend in (computational) neuroscience is to use Deep Learning to model biological brain regions and functions. This is motivated by the fact that deep neural networks (DNNs) combine brain inspired computational principles (Hassabis, Kumaran, Summerfield, & Botvinick, 2017; Khaligh-Razavi & Kriegeskorte, 2014; Kriegeskorte, 2015; Yamins & DiCarlo, 2016) with hitherto unseen effectiveness at solving perception tasks (He, Zhang, Ren, & Sun, 2016; Kriegeskorte, 2015; Krizhevsky, Sutskever, & Hinton, 2012; VanRullen, 2017). This renders DNNs well suited to uncover the kinds of representations and computations that may underlie complex, high-level functions of biological systems. As such, biologically plausible DNNs can be used as generative models to formulate new hypothesis about brain functionality (Cichy & Kaiser, 2019; Hassabis et al., 2017; Kriegeskorte & Douglas, 2018; Thompson, Bengio, Formisano, & Schönwiesner, 2018). For example, Sussillo, Churchland, Kaufman, and Shenoy (2015) used a recurrent neural network to find a naturalistic solution for the production of muscle activity related to reaching movements carried out by monkeys. Analysis of the trained network revealed that it learned to produce low-dimensional oscillator dynamics that generate multiphasic commands. This solution was predictive of both single-neuron and population-level activity observed in behaving monkeys. Furthermore, DNNs can be used to test hypotheses in neuroscience in silico by training them on ecologically relevant tasks and exposing them to stimuli used in neuroscientific experimentation. Wenliang and Seitz (2018) have, for example, used convolutional neural networks (CNNs) to study plasticity resulting from visual perceptual learning. The DNN reproduced key behavioral results and fulfilled predictions of existing specificity and plasticity theories regarding tuning changes in neurons of early primate visual areas. As another example, Kubilius, Bracci, and de Beeck (2016) exposed CNNs to visual stimuli to compare perceptual shape sensitivity between CNNs and humans. The authors hypothesized that a CNN’s success in capturing important aspects of human perception is due to a human-like representation of object shape. When trained for generic object recognition, the CNNs in this study learned physically correct object categories and demonstrated human shape sensitivity on several behavioral and neural stimulus sets.

Despite these successes, it is imperative to further improve the biological plausibility of DNNs in order for them to serve as sound models of neural systems (Hassabis et al., 2017; Thompson et al., 2018). For instance, while convolutional layers in CNNs exhibit some properties of early visual areas such as retinotopy and orientation tuning, they also lack properties found in early biological visual areas such as cortical magnification. This is the observation that, in early visual areas, the number of neurons dedicated to processing the visual field drops off with increasing eccentricity (Cohen, 2011; Daniel & Whitteridge, 1961). CNNs also lack a relationship between spatial frequency preference and eccentricity. Such a relationship has been observed in primate visual cortex, where small receptive fields exhibiting high spatial frequency preferences are concentrated foveally, whereas, with increasing eccentricity, receptive fields get progressively larger and tuned towards lower spatial frequencies (Broderick, Simoncelli, & Winawer, 2021; Dumoulin & Wandell, 2008; Sasaki et al., 2001; Smith, Singh, Williams, & Greenlee, 2001). Lastly, in contrast to CNNs, orientation tuning in the early visual cortex exhibits radial bias; the finding that radially oriented lines (and gratings) elicit a larger neuronal response than other orientations (Freeman, Brouwer, Heeger, & Merriam, 2011; Sasaki et al., 2006; Westheimer, 2003).

The aforementioned properties might be present due to inherent organizational principles of biological vision systems that are absent from CNNs. For example, it is assumed that cortical magnification is primarily explained by the decrease in retinal ganglion cell density with increasing eccentricity (Drasdo, 1977; Kwon & Liu, 2019; Wässle, Grünert, Röhrenbeck, & Boycott, 1989, 1990). The distribution of retinal ganglion cells exhibits a densely distributed central region, with an incrementally decreasing density as eccentricity increases (Curcio & Allen, 1990; Kwon & Liu, 2019; Watson, 2014). This results in a high sampling of the central visual field and low sampling of the periphery. When normalizing the distance between the cells (arranging them in a regular grid), the representational output resembles a wheelbarrow distortion of the retinal image in which the periphery is compressed while the fovea is stretched. A visual schematic of this effect can be seen in Figure 1.

**Figure 1:**
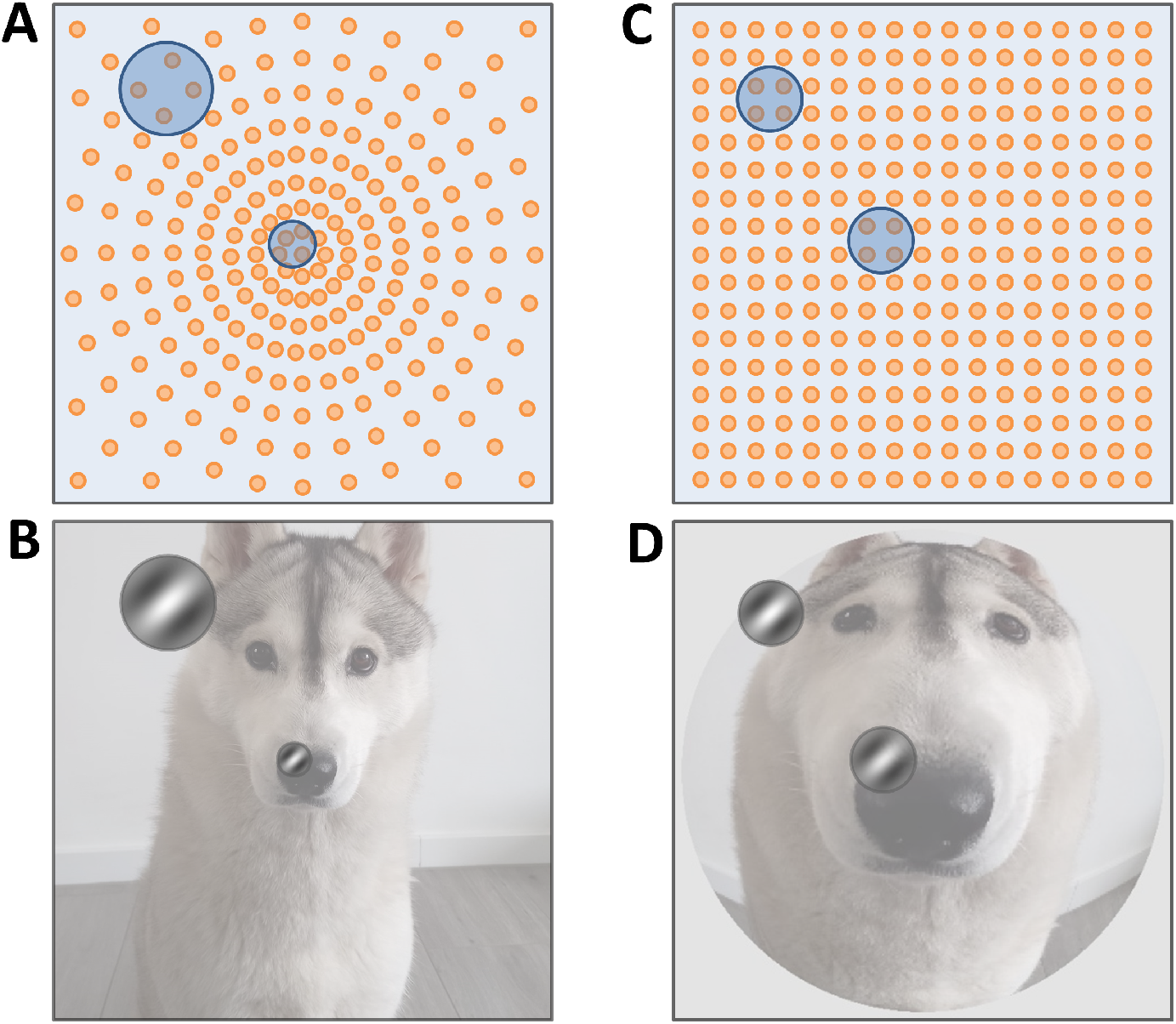
Schematic representation of retinal ganglion cell sampling. **A**, non-uniform distribution of ganglion cells. Spacing between cells increases as a function of eccentricity. **B**, differences in V1 receptive field sizes and spatial frequency tuning caused by non-uniform ganglion cell distribution. **C**, normalized ganglion cell distribution such that cells are equally spaced. **D**, distorted representation of the visual field as a result of equal spacing. Foveal areas are overrepresented compared to peripheral areas, whereas V1 receptive fields are now equal in size and spatial frequency preference.

Non-uniform sampling in the retina may not only give rise to cortical magnification but also to other eccentricity dependent properties. For example, under the assumption that the convergence rate of retinal ganglion cells to V1 cells is uniform across the visual field (as suggested by Kwon & Liu, 2019), eccentricity dependent differences in receptive field sizes in striatal areas may also be caused by the non-uniform retinal ganglion cell sampling (see again Figure 1 for illustration). Specifically, having fewer retinal ganglion cells at higher eccentricities but the same number of retinal ganglion cells converging to one V1 cell, V1 receptive fields will cover a larger portion of the visual field. Furthermore, the preference for low spatial frequencies at high eccentricity may be explained by the fact that high frequencies are compromised by the compression caused by low retinal ganglion cell density. With the same amount of retinal ganglion cells having to cover a larger part of the visual field, resulting in high distances between sampling points, the highest detectable frequency will decrease. Additionally, the wheelbarrow-like distortion resulting from retinal ganglion cell sampling does not affect radial lines, whereas it bends lines of other orientations. Since Gabor-like filters respond maximally to straight lines, the radial bias observed in biological vision systems (Freeman et al., 2011; Sasaki et al., 2006; Westheimer, 2003) may thus also be caused by the retinal ganglion cell sampling.

We utilize CNNs to test whether retinal ganglion cell processing combined with fixed convergence (i.e., location invariant receptive field size) can give rise to cortical magnification and increasing receptive field size as well as decreasing spatial frequency preference as a function of eccentricity. Furthermore, we test whether radial bias can result from retinal ganglion cell density combined with location invariant receptive field size and orientation tuning. Due to weight sharing, receptive fields in CNNs (filters) are location invariant and thus optimally suited for our purposes. We introduce a retinal sampling layer (RSL) which resamples images according to retinal ganglion cell density before feeding them into a shallow CNN. We train the resulting RSL-CNN on a simple orientation classification task, and subsequently compare the performance and emergent properties of the RSL-CNN to a reference CNN. We hypothesize that while introducing an RSL will not significantly impact classification performance, the RSL will introduce the previously mentioned characteristic properties found in the visual system. This would simultaneously lend credence to the conjecture that these properties emerge in the visual systems due to retinal ganglion cell density and render CNNs more biologically plausible.

## 2 Methods

### 2.1 Retinal Sampling Layer

The RSL takes a regularly sampled square image as input (non-square images will be zero-padded along the smaller dimension) and outputs a new, irregularly sampled square image. Irregular sampling in the output image corresponds to the irregular placement of ganglion cells in the retina. In short, the resampling operation converts between visual field coordinates and ganglion cell coordinates. Since the irregular placement of ganglion cells in the retina is eccentricity dependent and does not affect polar angles, it is convenient to use Polar coordinate systems for this conversion. Resampling is then essentially the effect of converting visual field radii (*r_vf_*) to ganglion cell radii (*r_gc_*); i.e. the number of ganglion cells in the interval [0, *r_vf_*]. We can obtain the number of ganglion cells by integrating a function relating ganglion cell density to eccentricity (visual field radius). For this, we use an empirically derived (Watson, 2014) ganglion cell density function:

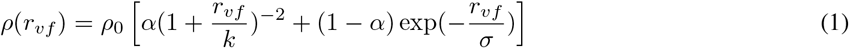

where *ρ*_0_ is the density at *r_vf_* = 0, *k* is the eccentricity at which density is reduced by a factor of four, *σ* is the scale factor of the exponential and *α* is the weighting of the first term. Parameter definition and values are shown in Table 1. The ganglion cell radius corresponding to a given visual field radius is then given by:

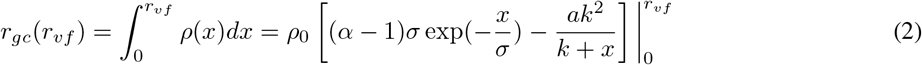

**Table 1.** Equation Parameters

| Symbol | Meaning | Value |
| --- | --- | --- |
| $\rho_0$ | ganglion cell density at $r_{vf} = 0$ | 33162 |
| $r_{vf}$ | Visual field radius in degrees | 20° |
| $k$ | $r_{vf}$ at which density is reduced by a factor of four | 1.05 |
| $\sigma$ | Scale factor of the exponential | 22 |
| $\alpha$ | Weighting parameter of the terms in equation 1 and 2 | 1 |

This relation allows one to move a pixel in the input image to its corresponding location in the output image. Specifically, after Cartesian pixel coordinates in the input image are (1) centered at zero by subtracting half of the image width (or height) from both the x- and y-coordinate and (2) converted to Polar coordinates, the visual field radius of a pixel is given by 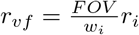, where *FOV* is the visual field of view covered by the input image, *w_i_* is the input image width and *r_i_* is the radius of the pixel in the input image space.

The field of view can be anywhere from 5° up to 100°, depending on the distance from which an image may be viewed. The radius of the pixel in the output image space is then 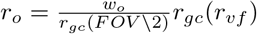, where *w_o_* is the output image width which can be freely chosen. Note that this latter conversion assumes that the convergence rate of retinal ganglion cells to V1 neurons is uniform across the visual field.

Such a forward resampling approach may lead to empty pixels in the output image. Therefore, it is preferable to employ an inverse resampling; i.e., to go through all pixels in the output image and query which pixels in the input image they correspond to. This renders it necessary to invert equation 2, which is complicated by the exponential term. However, since the exponential term is mainly necessary to capture ganglion cell density at very large eccentricities, we chose to omit it by setting *α* = 1.00. Please note that the contribution of the exponential term is generally small since *α* > 0.97 for all results reported by Watson (2014). Therefore, equation 2 becomes:

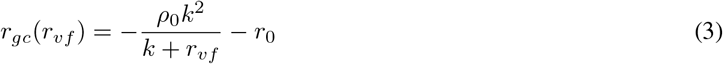

where 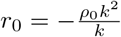 is the number of ganglion cells at zero eccentricity. The visual field radius is then given as a function of the ganglion cell radius

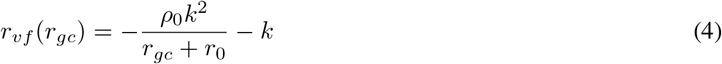

In most cases, pixel coordinates in the input image will not be integers. Therefore, we compute the intensity value of a pixel in the output image as the distance-weighted average over input image pixels whose coordinates are closest to the computed pixel coordinate. For fixed dimensions of input and output images and a fixed FOV, sampling can be performed by summarizing the relationship between pixels in the input and output image in a sparse weight matrix and multiplying this matrix with each of the vectorized RGB channels of the input image.

### 2.2 Neural Network and Training Data

A simple CNN architecture was used which consists of a single convolutional layer with rectified linear units (ReLU) leading into a softmax classification layer. Hyperparameter selection was based on a grid search method. The initial weights were initialized by sampling from a truncated normal distribution with a mean of zero and a standard deviation of 0.005. A filter size of 11-by-11 pixels with a stride of one is used. Additionally, zero-padding is applied such that the output size of the feature maps is equal to that of the input. Adadelta with standard hyperparameter values was used for network optimization (Zeiler, 2012). The only exception being that the initial learning rate was set to a value of 0.5. Training time was set to a standard 200 training epochs. Two versions of the network were used: a regular CNN, and the RSL-CNN in which the RSL precedes the first convolutional layer. The goal of the network was to classify the spatial orientation of sinusoidal gratings and Gabor patches. As CNN training usually results in Gabor-like filters, and since it is known that Gabor filters can be found in early visual areas, we want to investigate how the RSL affects the appearance of the filters. We varied the number of filters used for the convolution layer of both CNN and RSL-CNN networks. For every number of filters used for the convolutional layers, five training runs were performed. Using sparse categorical cross-entropy, we evaluated the trained models in terms of their classification accuracy on the training data.

Additionally, for the best performing model out of every five training runs, we investigated to which extent its filters are Gabor-like by calculating the correlation between each filter and a set of generated Gabor patches varying in orientation and size. We report the average and best performance of the models, and the averaged Gabor-likeness of the resulting filters in Appendix A. The number of filters used for further experimentation was chosen based on a hyperparameter optimization step, also described in Appendix A. All networks were constructed and tested using Keras with Tensorflow 2.4.0 on a NVIDIA Gigabyte RTX 3080 Aorus Master 10GB graphics card using NVIDIA CUDA 11.0 with cuDNN v8.2.1 support.

The training data consisted of 256-by-256 pixel sinusoidal gratings and Gabor patches. The sinusoidal gratings were created for 8 orientations (0, 22.5, 45, 67.5, 90, 112.5, 135, 157.5 degrees of polar angle), with 4 different spatial frequencies (0.0735, 0.147, 0.294, 0.5885 cycles per degree; c/deg), and 32 phase offsets (0 to 1.75*π*) per c/deg, per orientation. This resulted in 1024 grating stimuli with 128 examples per orientation. To generate the Gabor patches, 16 locations were selected for each orientation and for each spatial frequency. To prevent any Gabors from being (partially) presented outside of the visual field, the maximum radius was limited by 10 degrees of radial distance (eccentricity), minus 1.5 times the full width at half maximum (FWHM) of the Gaussian component of the Gabor. Four equidistant locations spanning the total image diameter were generated along four axes (0, 45, 90, 135 degrees of polar angle). This resulted in two low and two high eccentricity presentation locations for each Gabor type. For each location, both white and black-centered Gabors were created, leading to 32 stimuli per orientation, per frequency. In total, combining the gratings and Gabor patches resulted in 2048 images. Examples of these stimuli can be seen in Figure 2. For the RSL-CNN experiments, the aforementioned stimuli were initially created at a resolution of 2048-by-2048 pixels, and down sampled to 256-by-256 pixels using the RSL with a chosen FOV of 20°. These 20 degrees constitute the shortest distance between opposite stimulus borders. As generalization is not an issue here, all images were used for training, and overall training performance was used to evaluate model accuracy.

**Figure 2:**
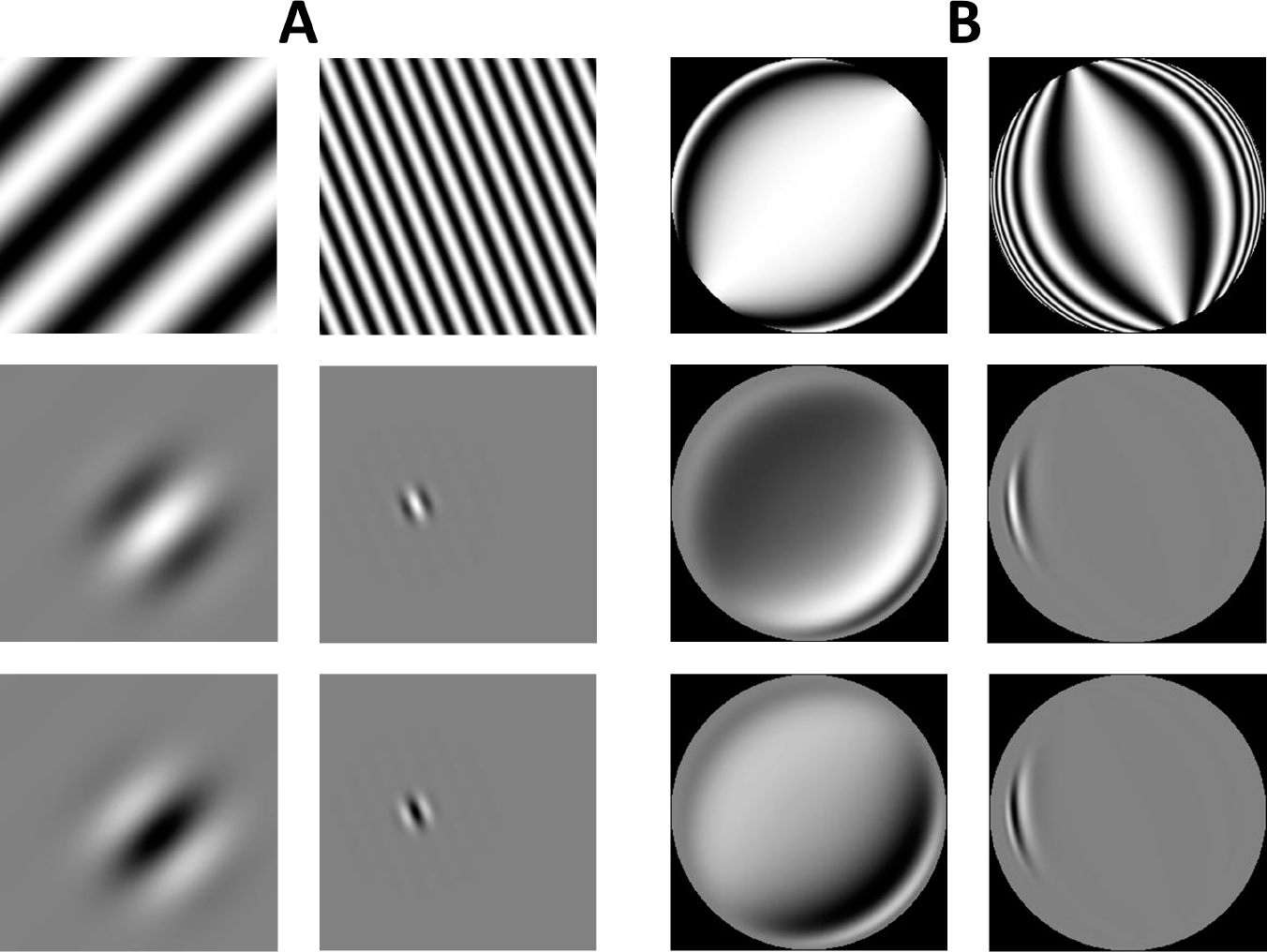
Stimuli used for training of the CNN (A), and RSL-CNN (B). The orientation indication signifies the orientation of the bar itself. **A**, exemplary sinusoidal gratings (top row), white-centered Gabor patches (middle row) and black-centered Gabor patches (bottom row). **B**, the same stimuli as in (A), after RSL processing.

### 2.3 Topography Analysis

#### 2.3.1 Retinotopy

We investigated the retinotopic organization of both the CNN and RSL-CNN by fitting location and size parameters of an isotropic Gaussian population receptive field (pRF) model (Dumoulin & Wandell, 2008) using a grid search method. To that end, we covered the input image with bar apertures for each of four orientations (0°, 45°, 90°, and 135°). The bar was 8 pixels wide and had a stride of 4 for each bar presentation. At each orientation-location combination, the bar aperture revealed each of the 1024 sinusoidal gratings used for training the network. Examples of these stimuli can be seen in Figure 3.

**Figure 3:**
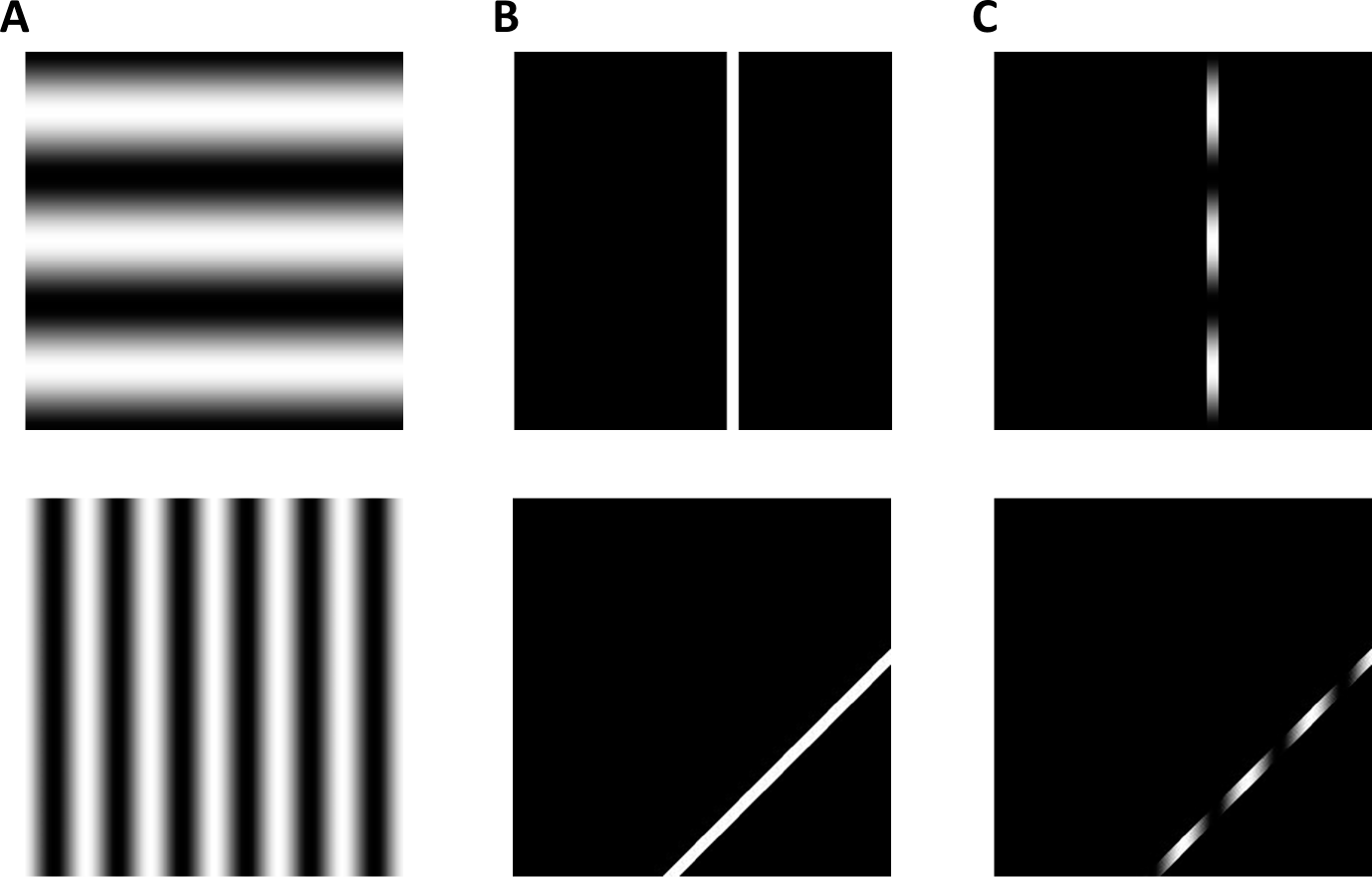
Two examples of stimuli used for the receptive field mapping. **A**, exemplary sinusoidal gratings. **B**, exemplary bar apertures. **C**, exemplary bar-masked sinusoidal gratings. For RSL-CNN studies, these stimuli were resampled using the RSL.

The layer activation at each visual field location (convolutional unit) to a single bar aperture was given by the average activity over all feature maps and gratings. An activation profile was created for every visual field location by recording the activations to all bar locations. We constructed candidate receptive fields as isotropic Gaussians at 16,384 uniformly distributed locations with sigma ranging from 0.03° to 1.2° in 15 steps for the grid search. We then obtained an activation profile for each receptive field candidate by computing the dot product between its vectorized Gaussian and each vectorized bar aperture. The pRF location and size of each convolutional unit were given by the location and sigma of the Gaussian whose activation profile correlated highest with the activation profile of that convolutional unit. A schematic overview of the adapted pRF mapping procedure can be seen in Figure 4.

**Figure 4:**
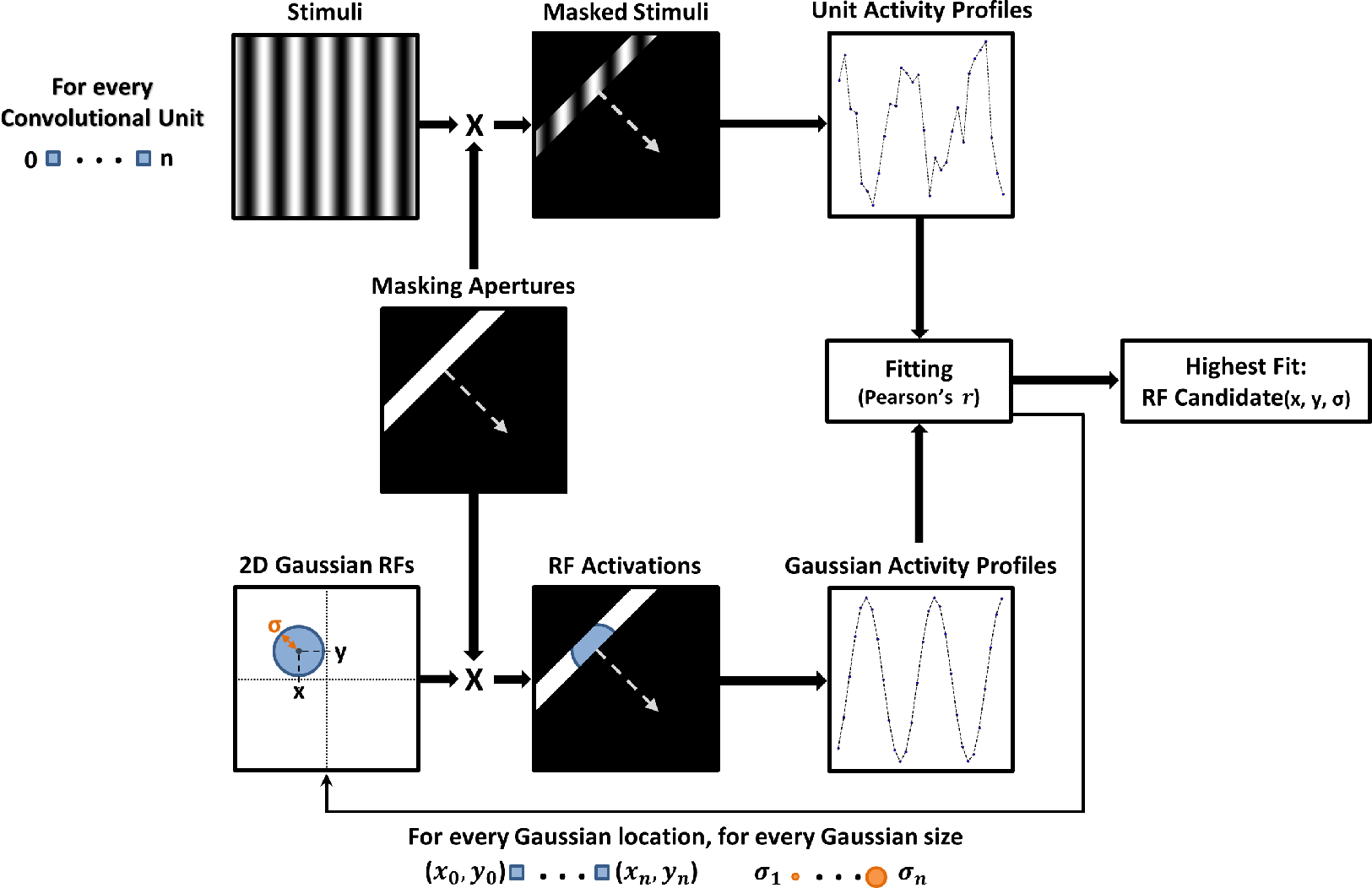
Schematic overview of the adapted pRF mapping method used for the convolutional layer in the neural networks. The top row demonstrates the creation of the neural network’s activity profiles, the bottom row shows the generation of the Gaussian activity profiles. The candidate receptive field for each convolutional unit is determined by the highest correlation coefficient of that unit’s activity profile and the activity profiles of all possible Gaussian receptive fields.

We report the eccentricity, polar angle, and size for each convolutional unit’s receptive field. Outlier removal for the eccentricity and polar angle values was based on the correlation between the activation profiles of the convolutional unit and the candidate receptive field. All cells with correlations which deviated from the mean by three standard deviations or more were removed. To describe the similarity between a model’s convolutional unit’s eccentricity and their corresponding eccentricity of the receptive field in the visual field, the weighted Jaccard similarity (*J_W_*) score was calculated. Additionally, to quantify the agreement between a model’s convolutional unit’s polar angle and their corresponding polar angle of the receptive field in the visual field it is vital to consider that this is a circular quantity.

As such, we decided to measure the congruence between polar angles using the Kuramoto order parameter. While the Kuramoto order parameter is typically employed to measure phase coherence among oscillatory time series, it is generally applicable to measure similarity between angles.

#### 2.3.2 Spatial-Frequency Preference

We investigated eccentricity-dependent preferences both for the CNN and RSL-CNN by exposing the trained models to achromatic sine-wave ring stimuli (sinrings) varying in eccentricity and spatial frequency (c.f. Henriksson, Nurminen, Hyvärinen, & Vanni, 2008; Mullen, Sakurai, & Chu, 2005). The rings were presented at five eccentricities (*e*; 1°, 2.8°, 4.7°, 6.6°, and 8.5°; image radius = 10°). The angular spatial frequencies were set at 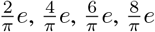, and 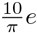 cycles per radian (cpr) to ensure equal spatial frequency content at each eccentricity. Examples of the stimuli are shown in Figure 5.

**Figure 5:**
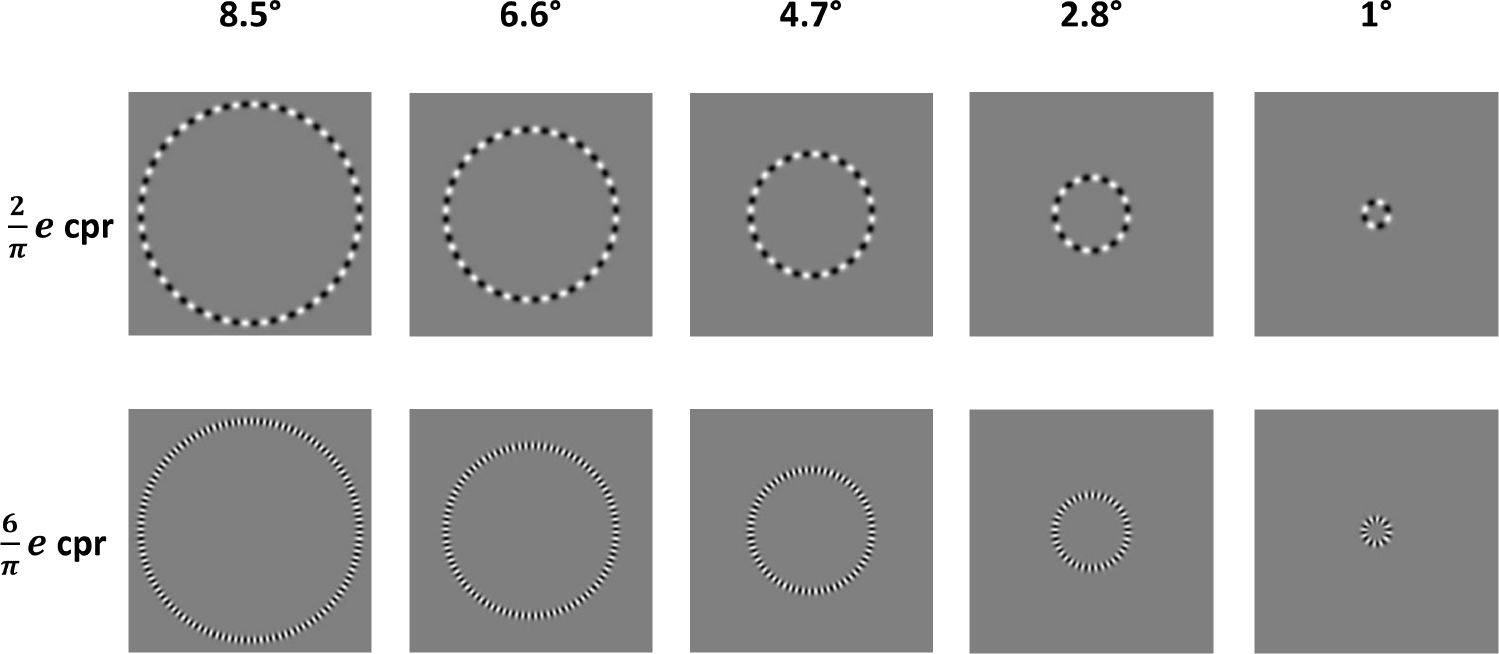
Examples of Sinring stimuli used for investigating eccentricity-dependent spatial frequency preferences. Each column displays a sinring at a different level of eccentricity (in degrees). Each row indicates a different angular frequency (in cycles per radian).

Per eccentricity level and for every spatial frequency, the sum of all convolution units of all output maps was calculated. The resulting value represents the network’s response to that eccentricity-frequency combination. Per eccentricity level, model responses to the different frequencies were normalized by dividing by the highest value. We considered the spatial frequency that yields the highest normalized response as the preferred spatial frequency at a given eccentricity level.

#### 2.3.3 Radial Bias

We investigated the presence of radial bias in the units of the convolutional layer for both the CNN and the RSL-CNN by exposing the networks to sinusoidal gratings of various orientations (0, 22.5, 45.0, 67.5, 90.0, 112.5, 135.0, and 157.5 degrees of polar angle), and spatial frequencies (0.1, 0.25, 0.5, 1, 2, 3, 4, 5, and 6 c/deg). Analysis of each network’s output was performed for every spatial frequency separately. First, for each stimulus, the output stimulus feature maps were averaged into one mean feature map. Second, all stimulus feature maps with the same stimulus orientation were averaged, resulting in orientation feature maps. Third, using a polar coordinate system on the orientation feature maps, each feature map was divided into 16 equally sized bins. Each bin covers exactly 45 degrees of polar angle and has 22.5 degree overlap with the next and previous bin. The mean polar angle of each bin indicates the direction of the bin. The directions of the bins were equal to the polar orientations of the stimuli. The mean activity of each bin was normalized, and taken as the average directional cell activity. A schematic overview of the method used to determine orientation bias in the networks is given in Figure 6.

**Figure 6:**
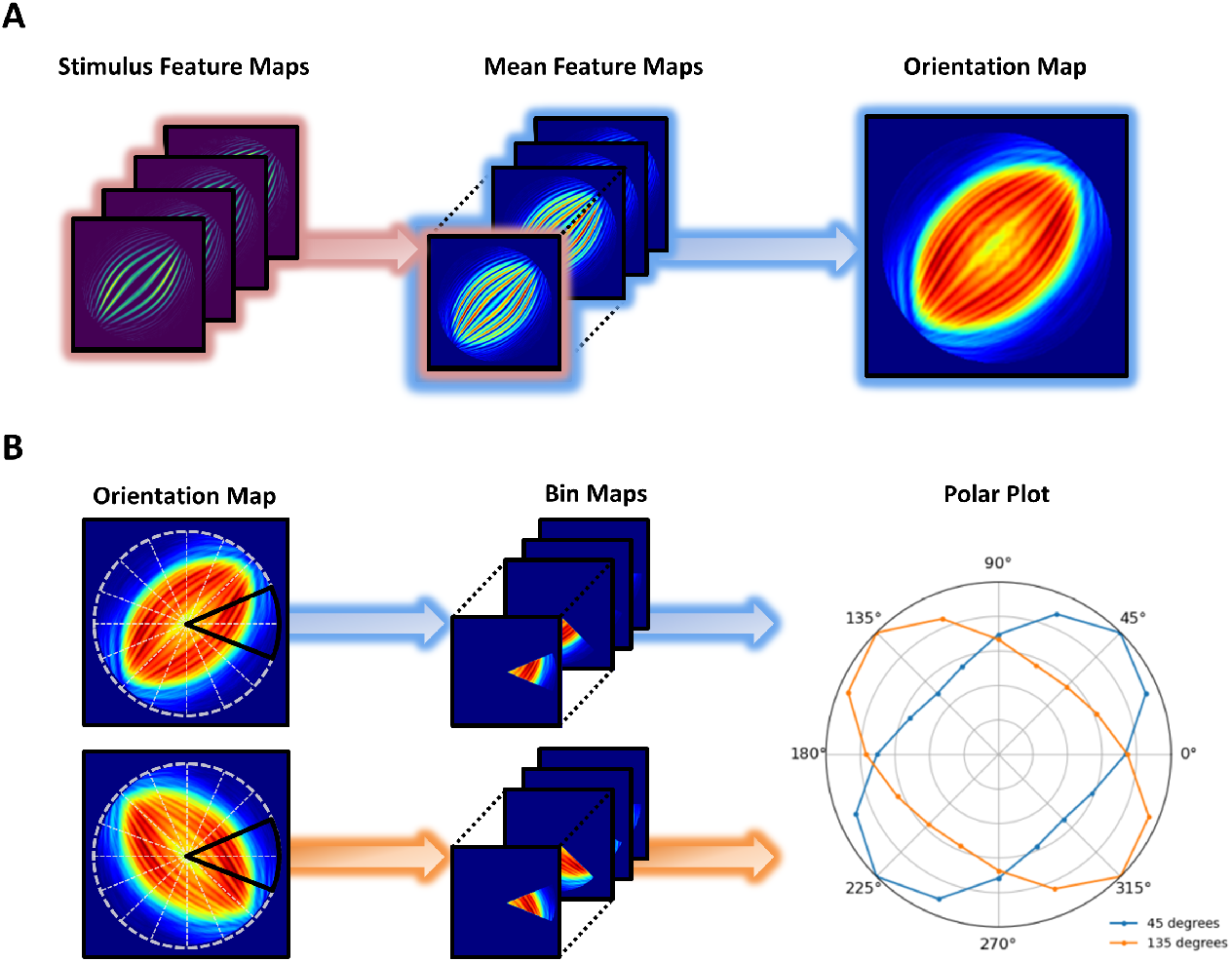
A schematic representation of the orientation mapping method. **A**, Feature maps for each stimulus are averaged into a mean feature map, resulting into one mean feature map for each stimulus. The mean feature maps of stimuli belonging to the same stimulus orientation are averaged into a single orientation map. **B**, Orientation maps are binned into 16 45-degree directional bins. The normalized activity per directional bin is then plotted into a polar plot.

As stimulus vignetting (Roth, Heeger, & Merriam, 2018), or stimulus edge effects (Carlson, 2014), may result in undesired non-stimulus-related activity, and since eccentricity is not of importance for the radial bias analyses, we excluded the outer 1 degree of eccentricity of the resulting feature maps before analyzing the layer activity. This resulted in the area outlined by the white dashed circle in left column of Figure 6B. The orientation bias for a given bar orientation is indicated by the bin with the highest response. To quantify radial bias we adapt the Kuramoto order parameter. Specifically, accounting for symmetry about the origin, radial bias is given by the average axial congruence 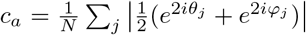 between the preferred orientation *θ_j_* and polar angle *φ_j_* (both in radians) of all units *j*. A value of 1 indicates perfect radial bias, while a value of 0 indicates an orthogonal bias (preferred orientation is orthogonal to polar angle).

## 3 Results

### 3.1 Retinotopy

By adapting the pRF mapping method to convolutional neural networks we obtained retinotopic maps for both the CNN and the RSL-CNN (see Figure 7). As expected, the polar angles in the RSL-CNN were unaffected by the distortion. As in the CNN, polar angles of network units in the RSL-CNN appeared to be in agreement with the visual angle of the image pixel to which they corresponded. Indeed, the congruency (*c*), as defined by the Kuramoto order parameter, between the polar angle of the convolutional unit of the network and the visual angle of corresponding image pixels was high for both networks with *c* = 1.00 and *c* = 0.99 for the CNN and RSL-CNN, respectively.

**Figure 7:**
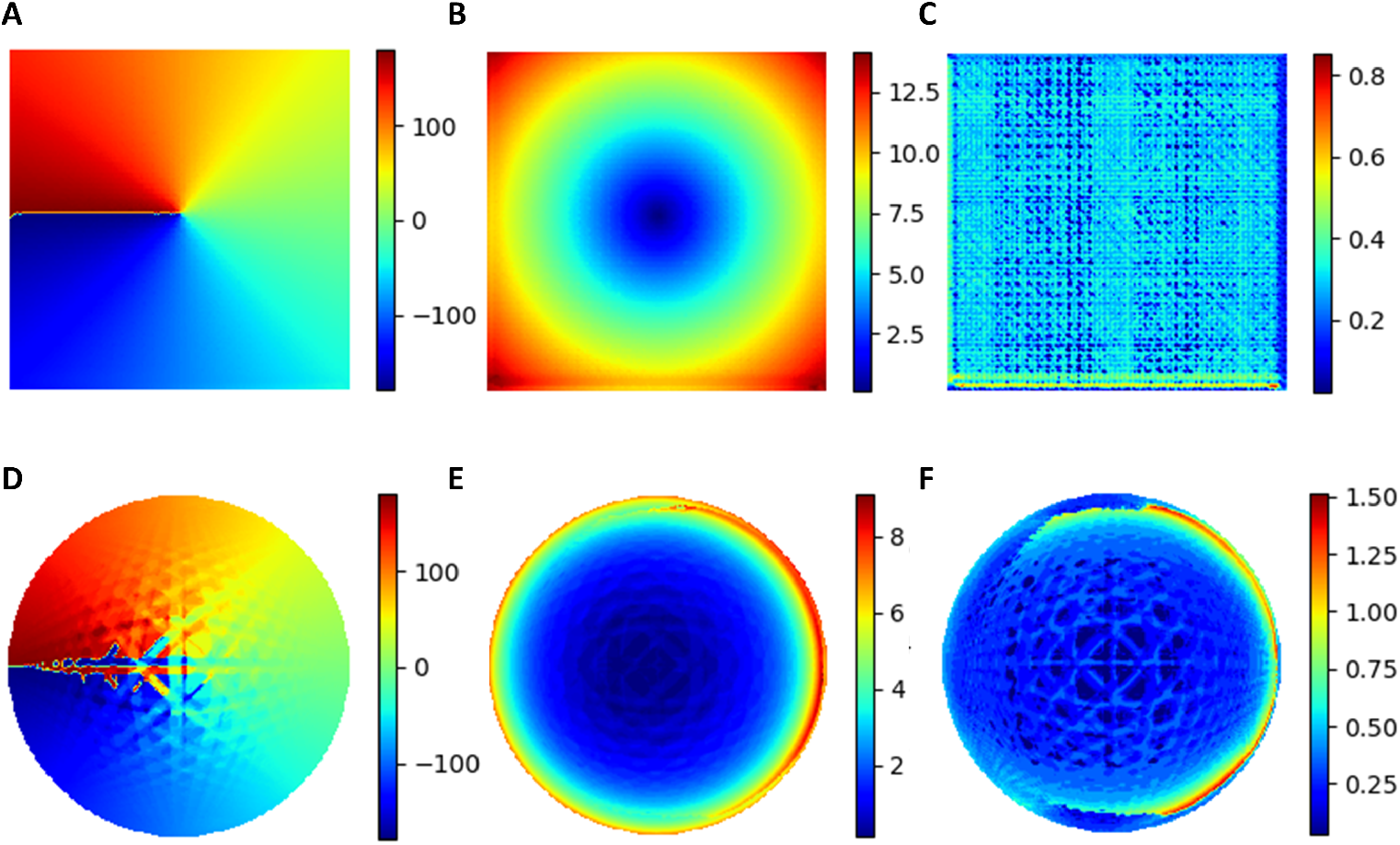
Retinotopic maps of convolutional layers. Each pixel in the maps reflects a convolutional unit in the corresponding convolutional layer. **A-C**, Polar angle, eccentricity, and receptive field size maps for the CNN. **D-F**, Polar angle, eccentricity, and receptive field size maps for the RSL-CNN.

For eccentricity, a different picture emerges. While for the CNN, eccentricities of the convolutional units were well aligned with the visual radius of corresponding image pixels (*J_W_* = 0.97), the same was not the case for the RSL-CNN (*J_W_* = 0.39). This discrepancy is due to cortical magnification which is apparent from the density of convolutional units dedicated to different eccentricities (see Figure 8a). This density was obtained by binning image pixels into rings whose width is equal to one degree of eccentricity, counting the number of convolutional units whose eccentricity falls within each bin and dividing this count by the area of the respective ring. For the CNN, which does not exhibit cortical magnification, density remained constant across eccentricities. For the RSL-CNN, however, density (*D*) was high at low eccentricities and dropped off exponentially as eccentricity (*E*) increases (*D* = 13737.133*E*^−1.012*E*^). In agreement with primate vision, a large region of the RSL-CNN’s convolutional layer was thus devoted to the visual field center.

**Figure 8:**
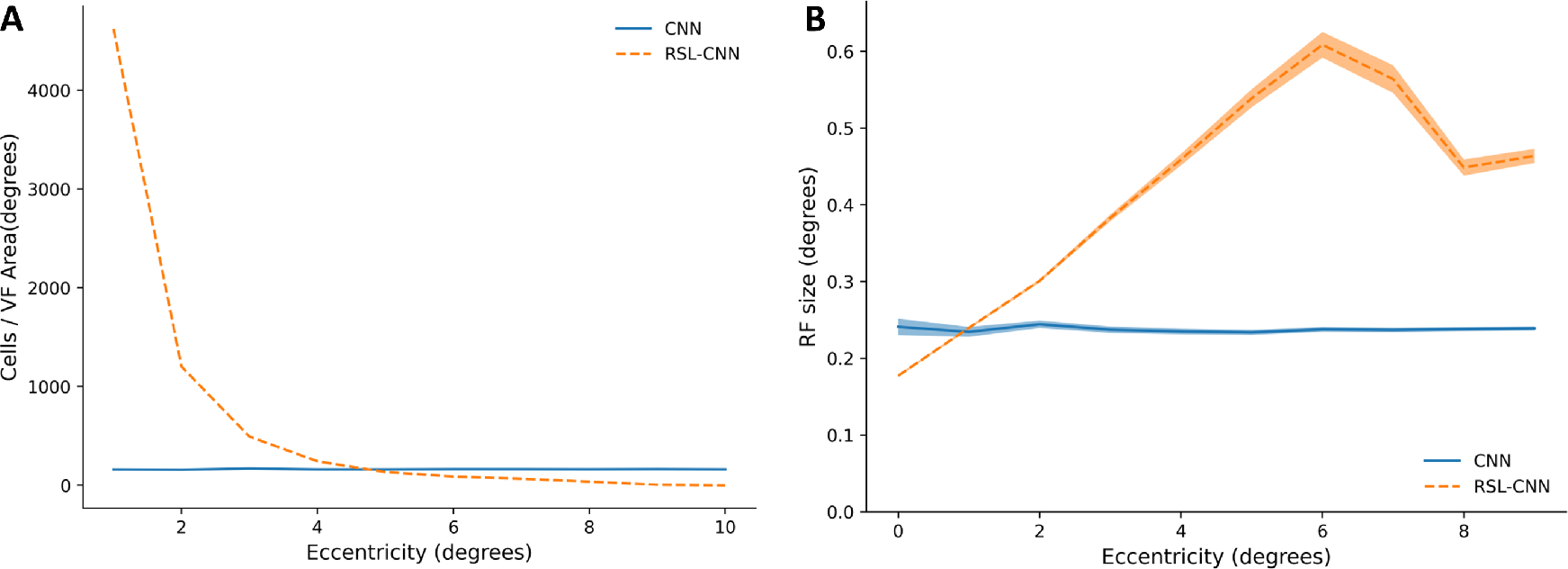
Properties of the convolutional layer for CNN and RSL-CNN. **A**, Number of convolutional units (cells) per degree of eccentricity, normalized for the visual field area. **B**, Average receptive field sizes per degree of eccentricity for a standard CNN and a RSL-CNN. The shaded area reflects the 1.96 standard error of measurement (SEM) scores.

Receptive field sizes in the RSL-CNN were also affected by the distortion caused by the retinal ganglion cell distribution. Receptive fields got larger with increasing eccentricity in the RSL-CNN (Figures 7f and 8b), whereas they remained constant for the CNN (Figures 6c and 7b). An asymmetry can be observed in Figure 6f, that cannot be explained by the Ganglion sampling algorithm, which is symmetric by nature. Furthermore, for the RSL-CNN’s a small left versus right asymmetry can be observed. The results of pRF mapping are affected by the chosen pRF parameters such as bar width and stride, and the amount, sizes, and position of the locations of the receptive field candidates. Additionally, the stimulus aperture vignetting (Roth et al., 2018) is not uniform but shows an activation bias to certain stimulus edges. This bias depends on the orientations of the convolutional filters: The closer a stimulus edge’s orientation is to the orientation of the available convolutional filters, the higher the stimulus edge effect will be. The previously mentioned pRF parameters, as well as the stimulus aperture vignetting, can have a distorting effect on the unit activation profiles. Nevertheless, while optimizing the previously mentioned parameters and reducing the stimulus aperture vignetting and edge effects could result in a better fit, it would not change the core observation that receptive field sizes increase with eccentricity.

Importantly, in accordance with primate visual systems, we observed that the relationship between receptive field size and eccentricity (E) was approximately linear and can be described as *RFsize_Model_* = 0.073*E* + .167. Note that the last three data points did not follow this trend. As the distortion of the visual field does not only result in progressively larger receptive fields, but also in radially compressed (i.e. aliased) and curved receptive field shapes, correlations with regular Gaussians may become less meaningful at higher eccentricities. Therefore, these data points were excluded when fitting the function.

### 3.2 Spatial-Frequency Preference

In the visual system, not only receptive field size is eccentricity-dependent but also spatial frequency preference (Broderick et al., 2021; Henriksson et al., 2008; Sasaki et al., 2001). We investigated whether such a relationship also emerges in the RSL-CNN but not the CNN by presenting sinring apertures revealing various spatial frequencies at a range of eccentricities to both neural networks (see Figure 9). Units in the CNN preferred a spatial frequency of 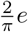 cpr irrespective of their eccentricity, with the normalized layer response dropping off as cpr increases. For the RSL-CNN, the spatial frequency preference shifted with eccentricity. More precisely, for units with the lowest eccentricity, the highest spatial frequency 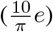 elicited the strongest overall network response. The lowest spatial frequency 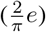 elicited the strongest overall network response for units with the highest eccentricity. In line with the primate visual system, the RSL-CNN thus showed a decrease in preferred spatial frequency as eccentricity increases.

**Figure 9:**
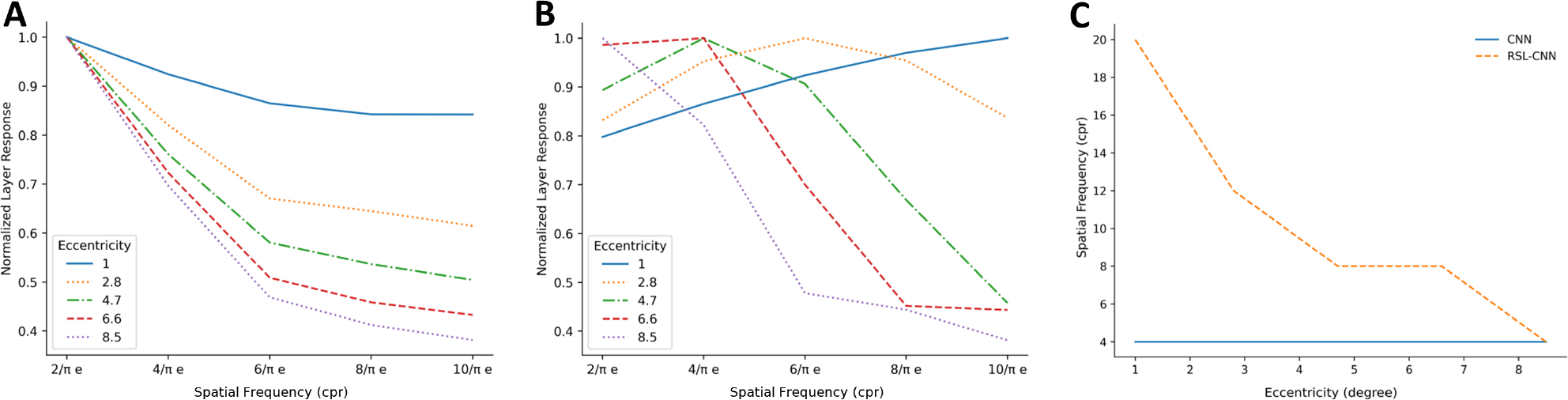
Eccentricity-dependent spatial frequency preference. **A**, Normalized averaged layer response of the CNN as a function of spatial frequency. Average responses observed at five eccentricities are presented separately. **B**, Same as A for the RSL-CNN. **C**, Spatial frequency preference as a function of eccentricity: CNN (blue) and RSL-CNN (orange).

**Figure 10:**
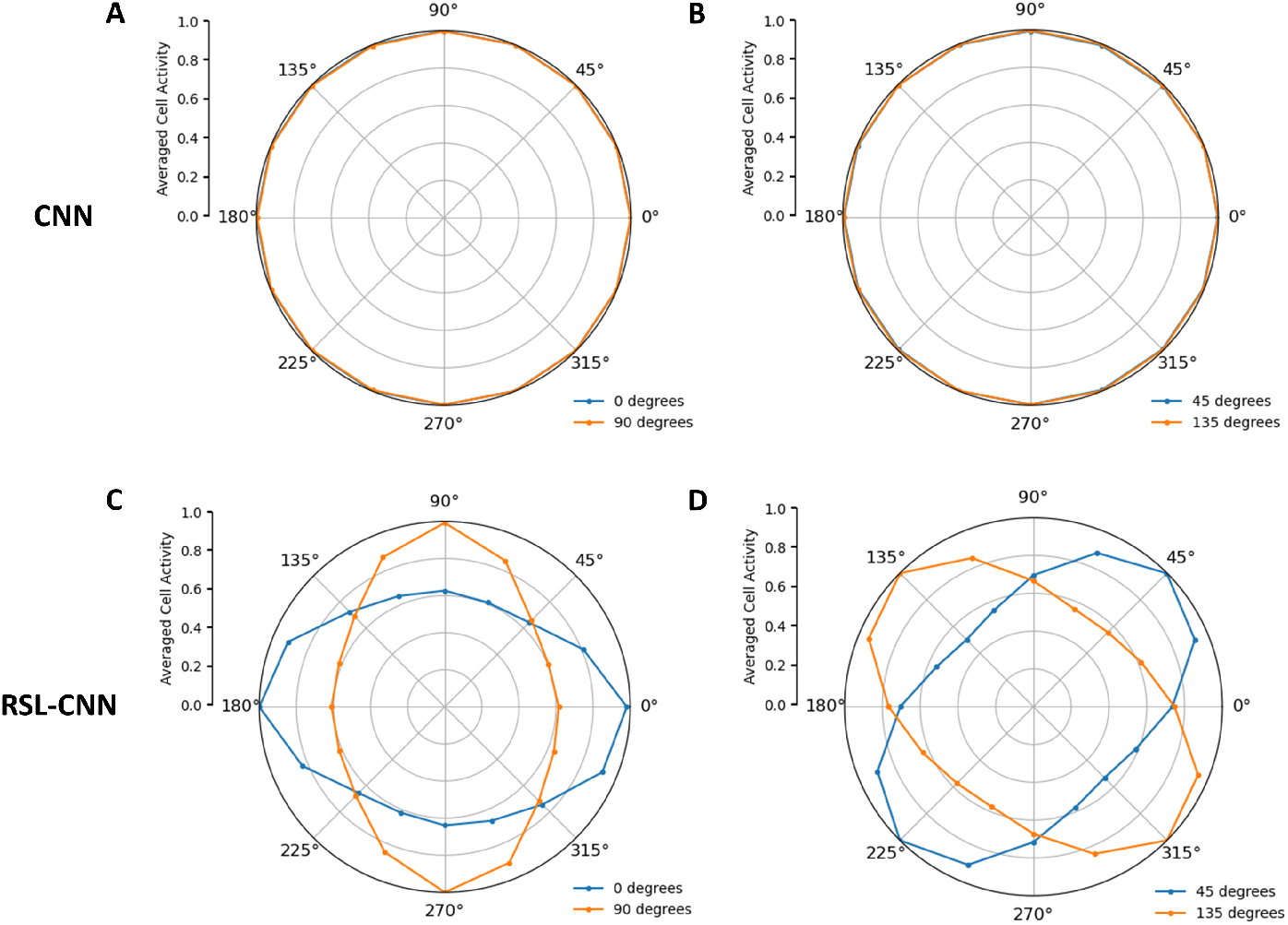
Radial bias plots for 4 c/deg. **A**, responses to 0° (orange) and 90° (blue) orientations of the CNN. **B**, responses to 45° (orange) and 135° (blue) orientations of the CNN. **C**, responses to 0° (orange) and 90° (blue) orientations of the RSL-CNN. **D**, responses to 45° (orange) and 135° (blue) orientations of the RSL-CNN.

### 3.3 Radial Bias

To investigate whether retinal ganglion cell sampling introduces radial bias, we presented sinusoidal gratings at four orientations (0°, 45°, 90° and 135°) to both networks. All gratings had a spatial frequency of 4 c/deg (c.f. Sasaki et al., 2006). As expected, the CNN showed no radial bias. The normalized directional cell activities are identical in every direction. The RSL-CNN, on the other hand, clearly exhibits radial bias. Responses were consistently strongest for units whose polar angle was coaxial with stimulus orientation.

## 4 Discussion

In the present study, we addressed the question whether non-uniform sampling of the visual field gives rise to some of the organizational principles of the visual system. To that end, we resampled images based on retinal ganglion cell distributions before feeding them into a simple convolutional neural network. Our results revealed that retinal ganglion sampling led the network to express human-like retinotopy, cortical magnification, eccentricity-spatial-frequency relationship, and radial bias. At the same time, image sampling only minimally impaired the network’s classification performance.

Population receptive field mapping applied to the RSL-CNN revealed that most of its receptive fields are located near the center of the visual field. Indeed, reminiscent of cortical magnification found in primate visual systems (e.g. Duncan & Boynton, 2003; Gattass, Gross, & Sandell, 1981), the number of convolutional units dedicated to a portion of the visual field decays exponentially with eccentricity. Furthermore, in combination with the non-uniform ganglion sampling of the visual field, the uniform convergence resulted in a biologically plausible eccentricity-related linear change of the receptive field sizes closely matching that observed in human V1. The receptive field-eccentricity slope of the RSL-CNN (0.073*E*) is similar to the slope of 0.072*E*, found in biological systems (*RFsize* = 0.072 · *RFeccentricity* + 0.017; Keliris, Li, Papanikolaou, Logothetis, & Smirnakis, 2019). A nonlinear distribution of retinal ganglion cells in conjunction with a fixed convergence rate from ganglion cells to the cortex can thus account for retinotopy as it implies that most V1 cells would receive input from a small central region and progressively fewer cells receive input from large peripheral regions. This lends further credence to the assumption that the convergence rate of retinal ganglion cells to V1 cells is indeed uniform across the visual field (as suggested by Kwon & Liu, 2019).

By comparing the intercept terms of the biological receptive field size (.017) versus the model receptive field size (.168; Keliris et al., 2019) as a function of eccentricity, it can be concluded that, given the 20 degrees FOV, receptive field sizes estimated within the network were smaller than those of biological receptive fields. It is important to note, however, that the effective receptive field size of the convolutional filter in the visual field depends on several factors: the resolution of the output image of the RSL, the FOV that is modeled by the RSL, and the receptive field size of the convolutional layer. For example, the effective coverage of the 11-by-11 pixels receptive field on a 256-by-256 pixel image that covers 20 degrees of visual angle is 0.86-by-0.86 degrees. If the size of RSL output is unchanged but covers 40 degrees of visual angle, the same receptive field (in pixels) would cover 1.72-by-1.72 degrees. Alternatively, if the resolution of the RSL output is changed to 512-by-512 pixels but still cover 20 degrees of visual angle, the same filter would cover 0.43-by-0.43 degrees of visual angle. Therefore, it is principally possible to match the receptive field size of the model to biological receptive field sizes by manipulating one, or several, of these parameters. Changing the effective visual field coverage of the convolutional filter may change the range of detectable spatial frequencies of a single filter, but its effect on the overall range of detectable spatial frequencies for the convolutional model cannot be inferred directly. Because of the nature of the convolutional process, the overall range of detectable spatial frequencies also depends on other factors such as filter stride. The discrepancy between the experimental results of the CNN and RSL-CNN demonstrates that the main findings are not dependent on the convolutional layer itself, but rather on the distortion of the visual field introduced by the RSL. Therefore, while the effective visual field coverage of the filters may change various details of our experimental outcomes, it would have no effect on the overall findings and conclusions.

Additionally, the estimated receptive field sizes for convolutional units seemed to be larger on the right half of the visual field than the left. As the retinal ganglion cell sampling is uniform in all angular directions, the cause for these deviating results cannot be the Ganglion sampling method by itself. A possible explanation for these deviating results is an imperfect pRF mapping method. An alternative explanation for the asymmetrical pRF size estimation may stem from the asymmetric zero-padding procedure of TensorFlow, the stimulus vignetting (Roth et al., 2018) and stimulus edge effects. Fine-tuning these methods further might refine our results but is not expected to lead to any changes in our general findings.

While retinal ganglion cell sampling is often believed to be the cause of cortical magnification (Drasdo, 1977; Kwon & Liu, 2019; Wässle et al., 1989, 1990), an alternative explanation would be that foveal ganglion cells have a disproportionately large representation in early visual areas (Malpeli & Baker, 1975; Myerson, Manis, Miezin, & Allman, 1977; Silveira, Picanço-Diniz, Sampaio, & Oswaldo-Cruz, 1989). The difference between these two theories lies in assuming a uniform convergence rate between the retinal ganglion cells and V1. While different magnification factors for LGN and striate cortex are proposed in the literature (e.g. Malpeli & Baker, 1975), our experiment provides further evidence that magnification in subsequent layers, closely resembling that of cortical magnification, arises from modeling just the ganglion density. The similarity of the magnification factors could potentially be improved by including additional mechanisms. Candidates for these would be the curvature of the eye, photoreceptor density, lateral spread of information in the retina via bipolar and amacrine cells, the convergence rates from retinal cells to cells in the lateral geniculate nucleus (LGN), and from the LGN to V1. Additionally, depending on the impact that additional mechanisms will have on the retinal representation, receptive field sizes can be matched to biological receptive field sizes. Given these results, it is likely that the resampling of images used here does not exactly reflect the input to V1. However, the general effect of resampling, namely a wheelbarrow-like distortion where the image center gets stretched whereas the periphery gets compressed, would remain the same if all additional factors were addressed. Since this type of distortion rather than its exact details gives rise to the organizational principles we discuss, we expect our results to hold at least qualitatively if these additional factors are included.

The RSL-CNN further exhibited eccentricity-dependent spatial frequency preferences. Like in primate visual systems (Broderick et al., 2021; Henriksson et al., 2008; Sasaki et al., 2001), the RSL-CNN exhibits a negative relationship between eccentricity and spatial frequency preference. This can be accounted for by higher spacing between retinal ganglion cells which results in a lower sampling rate at higher eccentricities, and hence, a compression of the input. In contrast, because of a high foveal ganglion cell density, low eccentricity input is forwarded to subsequent brain areas as an expanded representation. This increases the effective spatial frequency of stimuli presented at high eccentricities and reduces the effective spatial frequency of stimuli presented at low eccentricities. Applying a filter with a fixed orientation and spatial frequency tuning to the resampled image results in the filter preferring high spatial frequencies foveally, and low spatial frequencies in the periphery. In other words, a fixed filter’s preferred spatial frequency in retinal or cortical space implies decreasing spatial frequency preference as a function of eccentricity in the visual field.

Our findings indicate that the primate visual system may similarly exhibit receptive fields with constant effective spatial frequency tuning: Like in the RSL-CNN, in the primate visual system the receptive field size grows linearly as eccentricity increases (Dumoulin & Wandell, 2008; Smith et al., 2001), and the preferred spatial frequency grows linearly with eccentricity (Broderick et al., 2021; Henriksson et al., 2008; Sasaki et al., 2001). To translate our findings to primate visual systems, V1 cells with similar inherent orientation and spatial frequency selectivity should be present throughout the visual area, independent of the corresponding visual field location. This would imply that the empirically observed eccentricity-related tuning characteristics of visual cells (Broderick et al., 2021; Henriksson et al., 2008; Sasaki et al., 2001) are in fact not inherent tuning properties of the cells themselves. Rather, the observed differences in overall preferences may be caused by the non-linear retinal distortion of the visual field. In other words, a fixed convergence rate from ganglion cells to V1 cells, in combination with a uniform distribution of receptive fields with identical frequency and orientation preferences in early visual areas may account for these findings. This is in line with experimental findings that indicate that spatial frequency tuning is constant when expressed in cycles per millimeter cortex rather than per degree of visual angle (Rosa, Tweedale, & Elston, 2000; Virsu & Rovamo, 1979).

Finally, as in primate visual systems (Freeman et al., 2011; Sasaki et al., 2006; Westheimer, 2003), the RSL-CNN showed a radial bias effect for grating stimuli. Specifically, for sinusoidal gratings with 4 c/deg, the convolutional layer responses were highest along the radial axis parallel to the grating orientation. As expected, the radial bias can thus be accounted for by retinal ganglion cell distributions in conjunction with a constant converge rate of ganglion cells onto V1 cells, and constant effective spatial frequency and orientation tuning. Straight lines running perpendicular to radial directions are curved by the retinal ganglion cell sampling, while lines with a radial orientation are mostly unaffected. If identical Gabor-like filters are utilized throughout the visual field, the unaffected coaxial radial lines will activate these filters more than the curved lines that run perpendicular to the radial directions.

Because of weight-sharing, the aforementioned uniform application of identical filters across the visual input is ensured in a convolutional neural network. The assumption of weight sharing is based on the idea of the location invariance of statistically important features: a feature that is considered important in one location will also be important in a different location (Lecun & Bengio, 1995). Our results show that radial bias can occur when identical filters are applied uniformly across the visual input, without the need of location-specific filters. Therefore, as was suggested for the eccentricity-depended spatial frequency processing, it could be that the observed location-specific orientation sensitivity in primate visual systems (Freeman et al., 2011; Sasaki et al., 2006; Westheimer, 2003) is resulting from uniformly applying identical filters to retina-distorted visual input. Indeed, the visual system could have developed different versions of each filter at each location. For example, the brain could have curved Gabor filters at higher eccentricity to counteract the effect of the distortion. Our results indicate that this is not the case. Instead, a biological equivalent of weight-sharing, like having identical Gabor filters present throughout a visual area, in combination with the retina-distorted visual input, is sufficient for the network to display eccentricity-depending spatial frequency preferences and radial bias. This does not mean that only a single filter is available at a given location. Rather, as is the case with CNNs, different filter types can exist throughout the visual area.

The RSL-CNN exhibited lower classification performance compared to the original CNN. However, such a direct comparison is somewhat unjust. Where the CNN’s computing power is divided uniformly across the entire original input image, the ganglion sampling results in a high-resolution center representation, while peripheral information is represented in a compressed form. This directs the computational effectiveness of a CNN layer toward central areas of the input. As demonstrated in Apendix 1, this results in impaired classification performance at high eccentricities. The resulting loss of detailed peripheral information introduces a need for sequential integration of multiple aspects of the original image. In primate visual systems, this problem is solved by saccades and continuous integration of glimpses into a single mental representation. Our network architecture does not include such active sampling of the visual field. However, sequential integration of information extracted from a sequence of glimpses has been successfully addressed in the context of convolutional neural networks in the form of recurrent attention models (RAM; Mnih, Heess, Graves, & Kavukcuoglu, 2014). These models extract a partial observation from a base image through a GLIMPSE sensor. Depending on the size and number of patches, this method can lead to an intermediate representation that is larger than the original image. Additionally, because each new patch covers the same area as the previous patch, this method leads to redundant extractions of the same image regions. We suggest that the RSL be used as a more biologically realistic alternative to extract and downsize the region of interest with just a single matrix multiplication. Additionally, Ganglion sampling can be regarded as a non-uniform downsampling method in which information in the central region is kept intact, while the information in peripheral areas is represented in a compressed manner. The RSL layer downscaled the 2048-by-2048 pixels inputs to the 256-by-256 pixels image that was forwarded to the convolutional layer. With the cost of one RSL layer, the same CNN network successfully handled significantly larger input images. While uniform downscaling can reduce large images to a smaller size, it comes with the cost of losing information uniformly throughout the entire image. The RSL preserves information in the center of the image, while compressing the information in the periphery. Combining this concept of downscaling with the potential for sequential integration of scene aspects, such as provided by RAM models, could potentially lead to a very economical method for handling large images, or even full environments. Future research will explore the potential benefits of ganglion-based downscaling and RSL-based RAM models.

In conclusion, our results suggest that the interplay of three simple principles, namely, non-uniform distribution of ganglion cells, a uniform convergence rate from the retina to the cortex, and weight sharing, gives rise to retinotopy, eccentricity-dependent spatial frequency preferences, and radial bias closely resembling those found in primate visual systems. These findings support the idea that retinal ganglion cell sampling of the photoreceptors in the retina contributes to these organizational and functional properties of early visual areas. Additionally, these results expand our understanding of the functional organization of the visual system and offer more biologically realistic CNNs that may prove beneficial in machine vision.

## Acknowledgement

This study has received funding from the European Union’s Horizon 2020 Framework Programme for Research and Innovation under the Specific Grant Agreement Nos. 785907 (Human Brain Project SGA2) and 945539 (Human Brain Project SGA3).

## Appendices

### A Network Performance and Filter-Gabor Correlations

We investigated the effect that the number of available filters in the convolutional layer had on the performance and Gabor similarity of the filters. For every number of filters, five models were trained. The results of the training sessions are displayed in Table A1. As can be seen in Figure A1a, the general ability of the RSL-CNN to classify stimulus orientations (*M* = 0.93, *SD* = 0.032) was slightly impaired compared to the CNN (*M* = 0.98, *SD* = 0.057). This is expected as the RSL layer introduces a center bias towards which the convolutional layer’s computational resources are devoted. Therefore, high eccentricity information is impaired. We investigate this later with two selected sample networks. Next, we selected the best performing model per filter amount, resulting from the five training runs. All best performing CNN models performed similarly, close to 100% accuracy. For the RSL-CNN, it can be seen that performance increases as the number of filters increase.

For all the best-performing models per filter number group, we inspected the Gabor similarity of the filters. The average correlation coefficients for every filter amount tested can be seen in Figure A1b. The average correlations were calculated by transforming the correlation coefficient for every network using Fisher’s Z-scores. The Gabor similarity of all the best performing RSL-CNN models (*r*(10) = .82) was only slightly lower than that of the CNN models (*r*(10) = .85). This would suggest that, for the current dataset, the visual field distortion introduced by the RSL hardly impairs the resulting filters.

For further experimentation, a combined average score for best performance, average performance, and Gabor-filters correlation for both CNN and RSL-CNN was calculated for each number of available filters. As a sample network, a convolutional layer with four filters was selected as its combined classification performance, both with and without RSL, and the Gabor-likeness of the filters was highest. The best performing CNN with four filters had a classification accuracy of 99.90% with an average filter-Gabor correlation of *r*= .94. The best RSL-CNN with four filters had a classification accuracy of 89.65% and an average filter-Gabor correlation of *r*= .90.

We inspected the nature of the classification errors for both types of networks. It was found that, for the RSL-CNN, all sinusoidal gratings were classified correctly, while only 77.34% of the Gabor patches were classified correctly. Especially Gabors presented at high eccentricity levels were incorrectly classified (59.38% accuracy) compared to higher accuracy for low eccentricity Gabors (95.31% accuracy). In contrast, the CNN accurately classified 99.41% of the low and 99.61% of the high eccentricity Gabors.

Because the ganglion sampling creates a non-uniform distortion of the visual field in which the amount of distortion depends on the level of eccentricity, having more filters available will allow the convolutional layer to adapt to the different levels of distortion. The correlation of the trained filters with the predefined Gabors decreases as the number of available filters increase. Having more filters available allows the network to have more specific extraction suitable for this specific dataset. While this leads to higher accuracy, the Gabor similarity of the filters decreases. In general, it can be concluded that the Retinal Sampling Layer only slightly impacts the Gabor similarity and the performance of the models. Additionally, impaired performance for the RSL-CNN seems to be eccentricity-based. When the eccentricity of the presented object is increased, the chance of erroneous classification increases.

**Figure A1:**
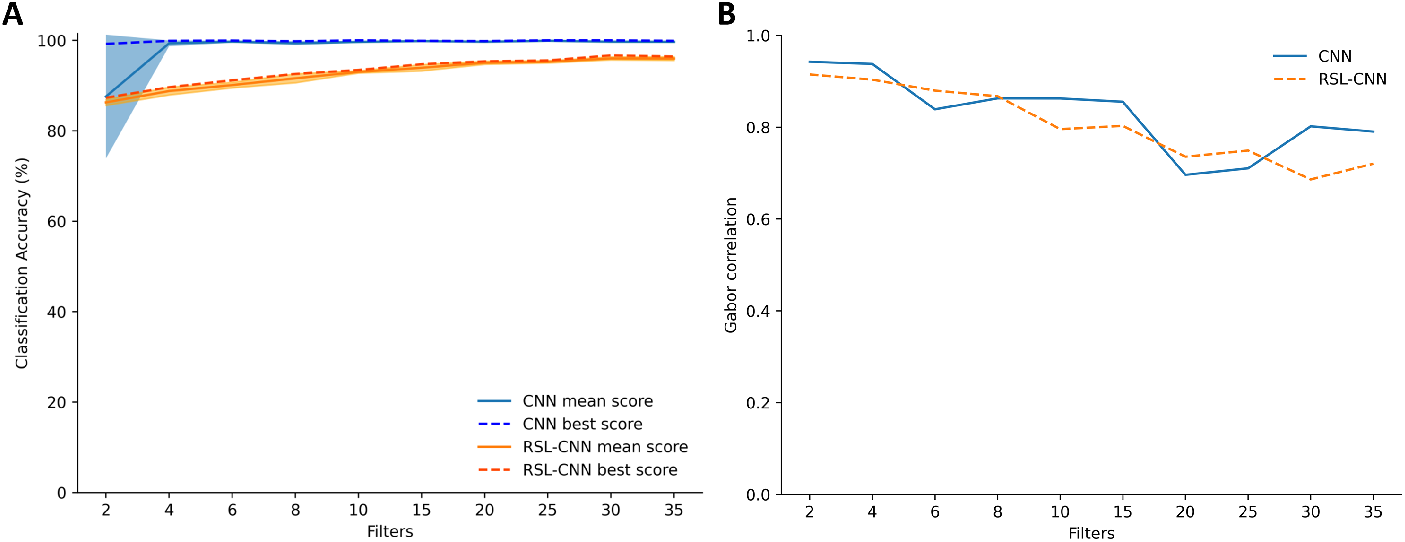
Model performance and Gabor correlations of the filters. **A**, Classification accuracy for the different amounts of filters used in the convolutional layer. **B**, Average Gabor correlations of the filters of the best performing models per filter group.

**Table A1.**
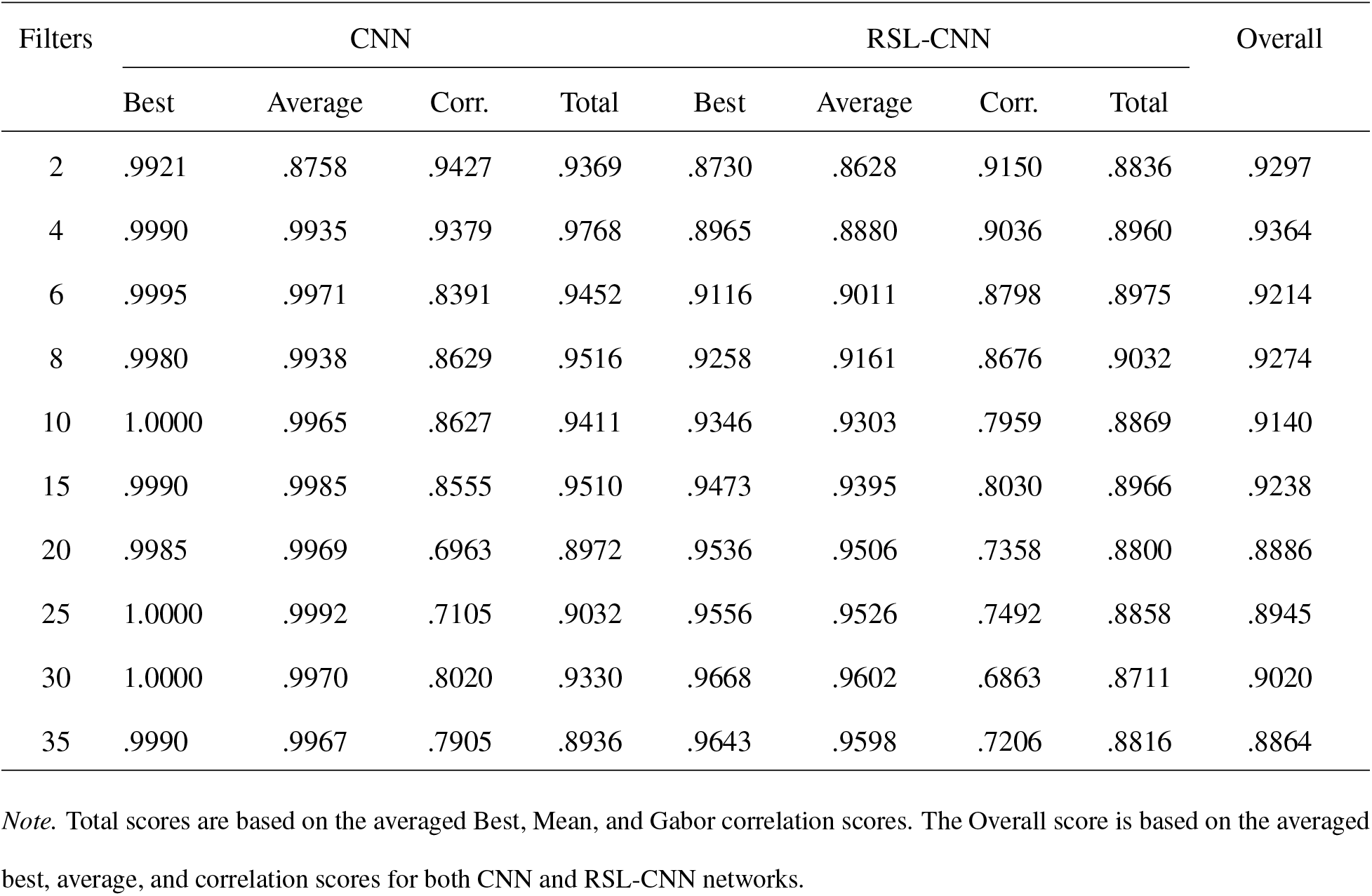
Performance scores and Gabor correlations for CNN and RSL-CNN training

| Filters | CNN |  |  |  | RSL-CNN |  |  |  | Overall |
| --- | --- | --- | --- | --- | --- | --- | --- | --- | --- |
|  | Best | Average | Corr. | Total | Best | Average | Corr. | Total |  |
| 2 | .9921 | .8758 | .9427 | .9369 | .8730 | .8628 | .9150 | .8836 | .9297 |
| 4 | .9990 | .9935 | .9379 | .9768 | .8965 | .8880 | .9036 | .8960 | .9364 |
| 6 | .9995 | .9971 | .8391 | .9452 | .9116 | .9011 | .8798 | .8975 | .9214 |
| 8 | .9980 | .9938 | .8629 | .9516 | .9258 | .9161 | .8676 | .9032 | .9274 |
| 10 | 1.0000 | .9965 | .8627 | .9411 | .9346 | .9303 | .7959 | .8869 | .9140 |
| 15 | .9990 | .9985 | .8555 | .9510 | .9473 | .9395 | .8030 | .8966 | .9238 |
| 20 | .9985 | .9969 | .6963 | .8972 | .9536 | .9506 | .7358 | .8800 | .8886 |
| 25 | 1.0000 | .9992 | .7105 | .9032 | .9556 | .9526 | .7492 | .8858 | .8945 |
| 30 | 1.0000 | .9970 | .8020 | .9330 | .9668 | .9602 | .6863 | .8711 | .9020 |
| 35 | .9990 | .9967 | .7905 | .8936 | .9643 | .9598 | .7206 | .8816 | .8864 |

*Note.* Total scores are based on the averaged Best, Mean, and Gabor correlation scores. The Overall score is based on the averaged best, average, and correlation scores for both CNN and RSL-CNN networks.

